## Supporting Information for "CryoEM of endogenous mammalian V-ATPase interacting with the TLDc protein mEAK-7"

[1]Molecular Medicine Program, The Hospital for Sick Children Research Institute, Toronto, ON M5G 0A4, Canada.
[2]Program in Cell Biology, The Hospital for Sick Children, Toronto, ON M5G 0A4, Canada.
[3]Department of Biochemistry, University of Toronto, Toronto, ON M5S 1A8, Canada.
[4]Physical and Theoretical Chemistry Laboratory, University of Oxford, Oxford, OX1 3QZ, UK.
[5]Kavli Institute for Nanoscience Discovery, University of Oxford, Oxford, OX1 3QU, UK.
[6]Lunenfeld-Tanenbaum Research Institute, Sinai Health System, Toronto, ON M5G 1X5, Canada.
[7]Department of Molecular Genetics, University of Toronto, Toronto, ON M5S 1A8, Canada.
[8]Department of Medical Biophysics, University of Toronto, Toronto, ON M5G 1L7, Canada.
^Current address: Department of Biological Sciences, National University of Singapore, 16 Science Drive 4, Singapore 117558, Singapore.
[&]Current address: Disease Intervention Technology Laboratory, Immunos, Agency for Science, Technology and Research (A*STAR), 8A Biomedical Grove, Singapore 138648, Singapore.

**Supporting information:**
- Two supplementary data Microsoft Excel spreadsheets
- One supplementary video
- Three supplementary tables
- Eight supplementary figures

**Supplementary Data 1. Proteins identified by mass spectrometry from each gel region in purified porcine kidney V-ATPase.**

**Supplementary Data 2. Mass spectrometry comparison of isoform content for different V-ATPase subunits.**

**Video 1. Illustration of mEAK-7 binding and unbinding V-ATPase.**

**Supplementary Table 1. Cryo-EM data collection and modeling statistics for V-ATPase alone**

| | State1 | State2 | State3 |
|---|---|---|---|
| **EM data collection / processing** | | | |
| Microscope | | FEI Titan Krios | |
| Voltage (kV) | | 300 | |
| Camera | | Falcon 4 | |
| Mode | | Counting | |
| Defocus mean ± std (µm) | | 1.7 ± 0.4 | |
| Exposure time (s) | | 9 | |
| Number of fractions | | 29 | |
| Exposure rate ($e^-$/pixel/s) | | 5 | |
| Total exposure ($e^-$/Å$^2$) | | 40 | |
| Pixel size (Å) | | 1.01867 | |
| Number of micrographs | | 5,451 | |
| Number of particles (after cleanup) | | 133,337 | |
| Number of particles (in final map) | 24,327 | 14,746 | 22,866 |
| Symmetry | C1 | C1 | C1 |
| Resolution (global) (Å) | 3.7 | 4.1 | 3.8 |
| Directional Resolution Range (Å) | 3.5-3.9 | 3.9-5.0 | 3.6-4.2 |
| Sphericity of 3DFSC | 0.92 | 0.74 | 0.89 |
| SCF Value* | 0.84 | 0.84 | 0.84 |
| Map B factor (Å$^2$) | -76 | -70 | -77 |
| EMDB ID | TBC | TBC | TBC |
| EMPIAR ID | TBC | TBC | TBC |
| **Model statistics** | | | |
| Residues | 8978 | 8978 | 8978 |
| Ligand | ADP | ADP | ADP |
| Map CC | 0.78 | 0.70 | 0.76 |
| RMSD [bonds] (Å) | 0.021 | 0.020 | 0.021 |
| RMSD [angles] (Å) | 1.724 | 1.802 | 1.777 |
| All-atom clashscore | 0.84 | 0.85 | 0.82 |
| Ramachandran plot | | | |
| Favored (%) | 97.74 | 97.68 | 97.76 |
| Allowed (%) | 2.01 | 2.06 | 2.02 |
| Outliers (%) | 0.25 | 0.26 | 0.21 |
| Rotamer outliers | 0.01 | 0.06 | 0.08 |
| C-β deviations | 0.06 | 0.07 | 0.04 |
| MolProbity Score | 0.82 | 0.83 | 0.81 |
| EM-Ringer Score | 1.26 | 0.42 | 1.05 |
| PDB ID | TBC | TBC | TBC |

*The sampling compensation factor (SCF) value is calculated as described (Baldwin and Lyumkis, 2020) and assumes that all orientations have been determined accurately.

**Supplementary Table 2. Cryo-EM data collection and modeling statistics for mEAK7-bound V-ATPase structures**

| **Dataset** | **V-ATPase + mEAK7** | | **V-ATPase + mEAK7ΔCterm** | | **V-ATPase + mEAK7 + ATP** | | **V-ATPase + mEAK7 + EDTA + EGTA** | **V-ATPase + mEAK7 + Calcium** |
|---|---|---|---|---|---|---|---|---|
| State | State 2 loosely bound mEAK7 | State 2 tightly bound mEAK7 | State 2 no mEAK7 | State 2 bound mEAK7 | State 2 no mEAK7 | State 2 bound mEAK7 | State2 bound mEAK7 | State2 bound mEAK7 |
| **EM data collection / processing** | | | | | | | | |
| Microscope | Titan Krios | | Titan Krios | | Titan Krios | | Titan Krios | Glacios |
| Voltage (kV) | 300 | | 300 | | 300 | | 300 | 200 |
| Camera | Falcon 4 | | Falcon 4 | | Falcon 4 | | Falcon 4 | Falcon 4 Selectris X |
| Mode | Counting | | Counting | | Counting | | Counting | Counting |
| Defocus mean ± std (μm) | 2.0 ± 0.4 | | 1.8 ± 0.4 | | 1.9 ± 0.5 | | 2.3 ± 0.5 | 0.8 ± 0.2 |
| Exposure time (s) | 17 | | 8 | | 11 | | 10 | 5 |
| Number of fractions | 29 | | 29 | | 29 | | 29 | EER |
| Exposure rate ($e^-$/pixel/s) | 4 | | 9 | | 8 | | 8 | 6 |
| Total exposure ($e^-$/Å$^2$) | 36 | | 37 | | 44 | | 41 | 40 |
| Pixel size (Å) | 1.3225 | | 1.3225 | | 1.3225 | | 1.3225 | 0.895 |
| Number of micrographs | 5,142 | | 2,013 | | 5,498 | | 2,468 | 3,029 |
| Number of particles (after cleanup) | 186,338 | | 107,515 | | 253,892 | | 102,191 | 123,373 |
| Number of particles (in final map) | 11,437 | 31,814 | 15,576 | 4,315 | 14,065 | 70,360 | 24,994 | 8,611 |
| Symmetry | C1 | C1 | C1 | C1 | C1 | C1 | C1 | C1 |
| Resolution (global) (Å) | 4.2 | 3.5 | 4.2 | 6.6 | 4.2 | 3.6 | 3.7 | 4.1 |
| Directional Resolution Range (Å) | 4.0-6.6 | 3.4-3.7 | 4.0-5.7 | 5.9-7.4 | 4.1-5.5 | 3.4-3.7 | 3.5-4.0 | 3.9-6.0 |
| Sphericity of 3DFSC | 0.72 | 0.89 | 0.82 | 0.95 | 0.82 | 0.97 | 0.91 | 0.70 |
| SCF Value* | 0.68 | 0.59 | 0.80 | 0.76 | 0.78 | 0.76 | 0.76 | 0.72 |
| Map B factor (Å$^2$) | -97 | -107 | -95 | -247 | -98 | -126 | -99 | -10 |
| EMDB ID | TBC | TBC | TBC | TBC | TBC | TBC | TBC | TBC |
| EMPIAR ID | TBC | TBC | TBC | TBC | TBC | TBC | TBC | TBC |

*The sampling compensation factor (SCF) value is calculated as described (Baldwin and Lyumkis, 2020) and assumes that all orientations have been determined accurately.

**Supplementary Table 3. Cryo-EM data collection and modeling statistics for Merged Data**

| | V-ATPase + mEAK7 (State 2 tightly bound mEAK7) & V-ATPase + mEAK7 + EDTA + EGTA (State2 bound mEAK7) |
|---|---|
| **EM processing** | |
| Number of particles (in final map) | 56,808 |
| Symmetry | C1 |
| Resolution (global) (Å) | 3.5 |
| Directional Resolution Range (Å) | 3.3-3.7 |
| Sphericity of 3DFSC | 0.97 |
| SCF Value* | 0.70 |
| Map B factor (Å$^2$) | -118 |
| EMDB ID | TBC |
| **Model statistics** | |
| Residues | 9348 |
| Ligand | - |
| Map CC | 0.82 |
| RMSD [bonds] (Å) | 0.016 |
| RMSD [angles] (Å) | 1.798 |
| All-atom clashscore | 2.33 |
| Ramachandran plot | |
| Favored (%) | 97.60 |
| Allowed (%) | 2.19 |
| Outliers (%) | 0.22 |
| Rotamer outliers | 0.61 |
| C-β deviations | 0.17 |
| MolProbity Score | 1.10 |
| EM-Ringer Score | 1.64 |
| PDB ID | TBC |

*The sampling compensation factor (SCF) value is calculated as described (Baldwin and Lyumkis, 2020) and assumes that all orientations have been determined accurately.

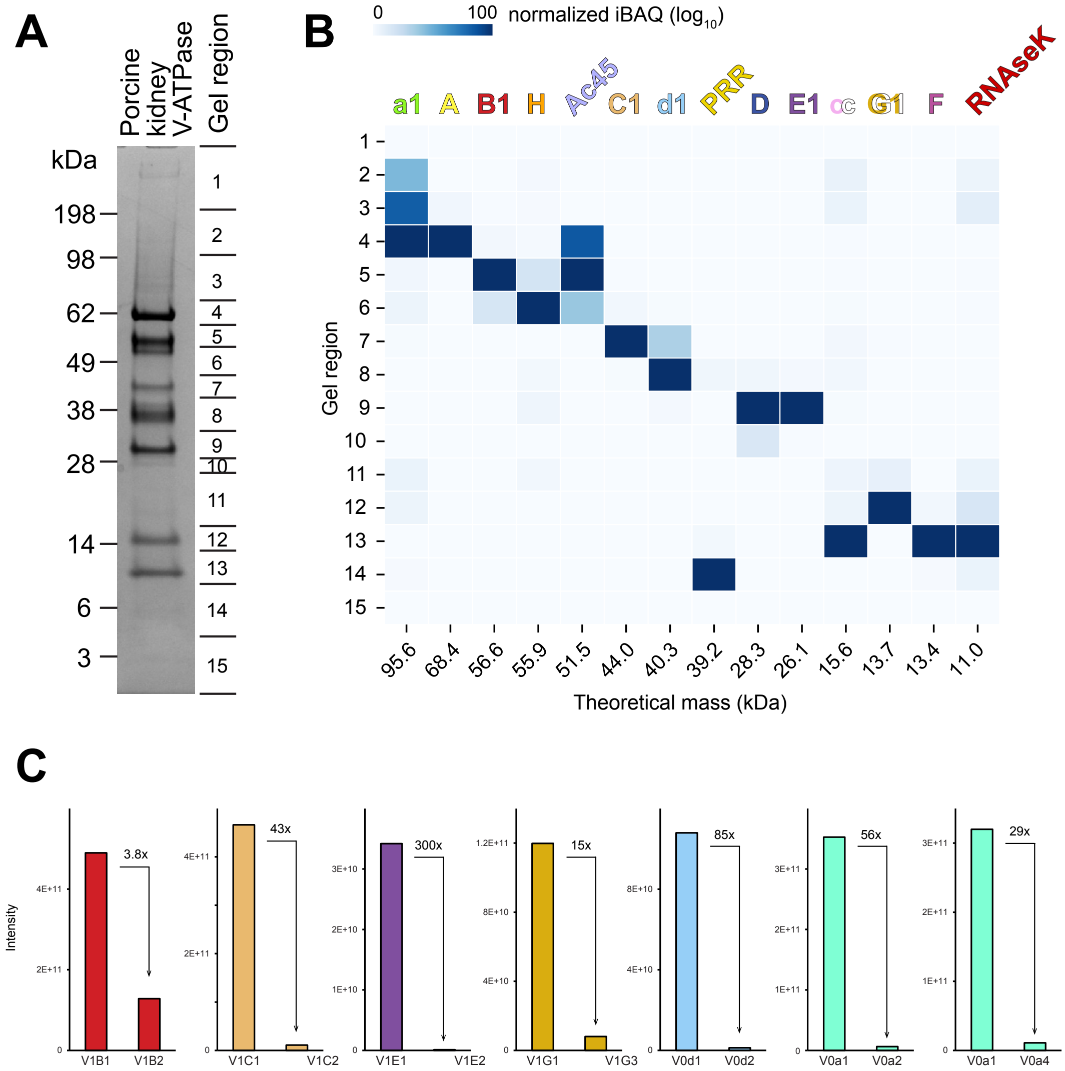


**Supplementary Figure 1. Mass spectrometry identification of subunit isoforms. A,** Gel regions subjected to mass spectrometry. **B,** Subunit isoforms identified in each gel region. The colour density represents the normalized $\log_{10}$(iBAQ score). **C,** Approximate abundance of different subunit isoforms (iBAQ score).

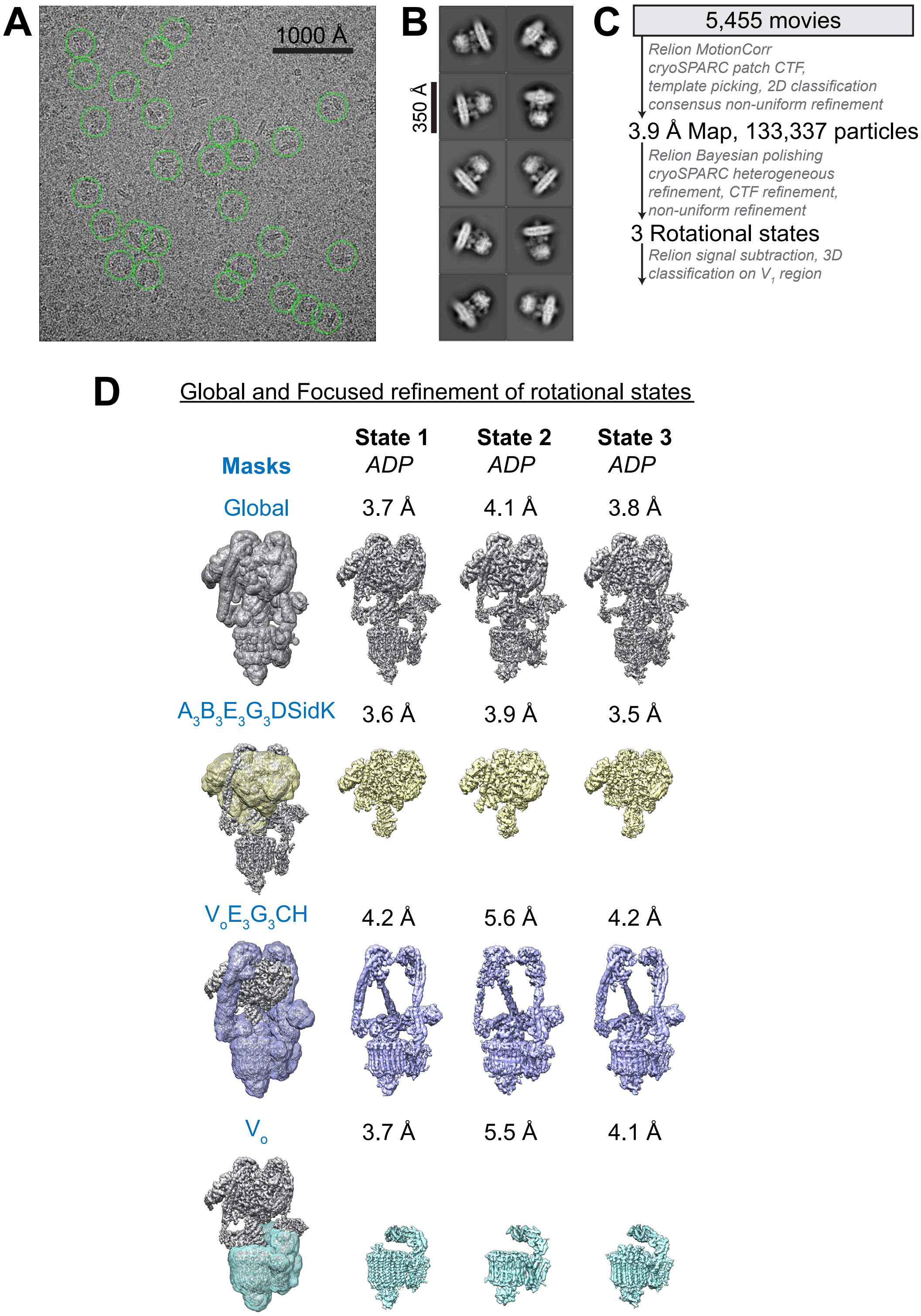


**Supplementary Figure 2. CryoEM workflow. A,** Example micrograph. **B,** Example class average images. **C,** Workflow used to identify different conformations of V-ATPase. **D,** Focused refinement scheme used to produe high-resolution maps.

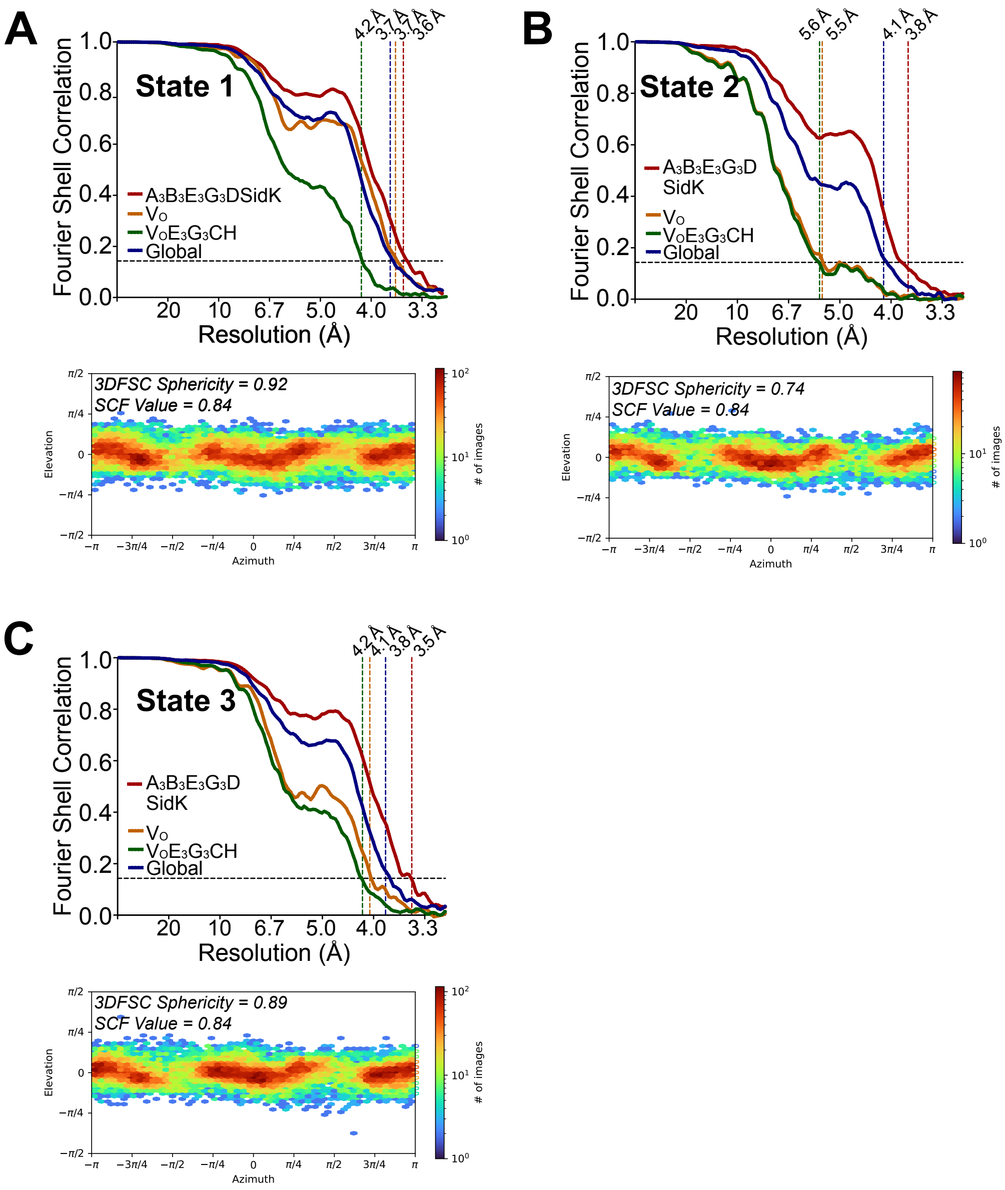


**Supplementary Figure 3. CryoEM map validation.** Fourier Shell Correlation curves following a gold-standard refinement and correction for masking, and orientation distribution plots, for rotational State 1 (**A**), State 2 (**B**), and State 3 (**C**).

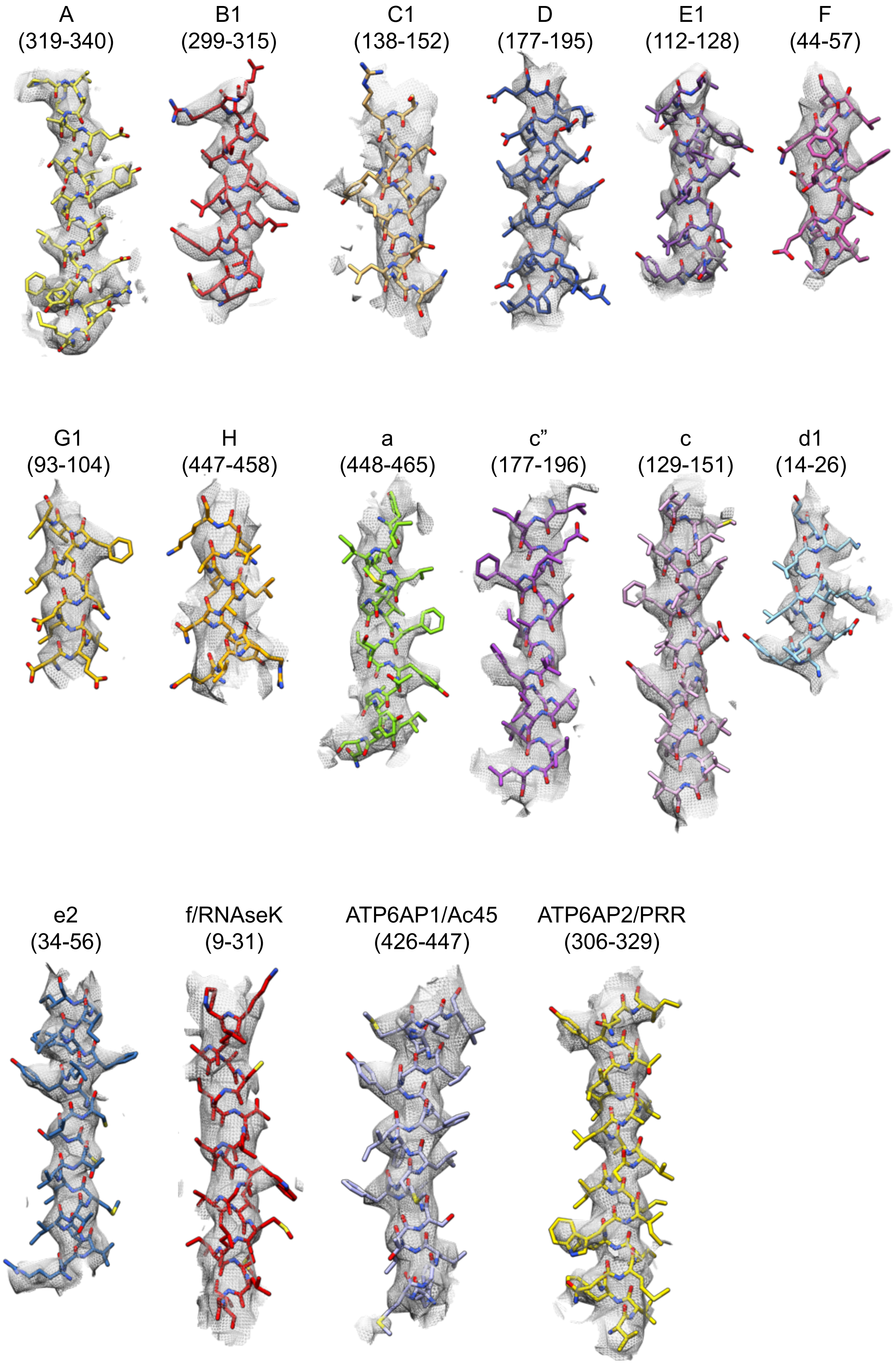


**Supplementary Figure 4. Examples of model-in-map fit.** Examples are shown from each of the different V-ATPase subunit types. Residue numbers shown are indicated in brackets.

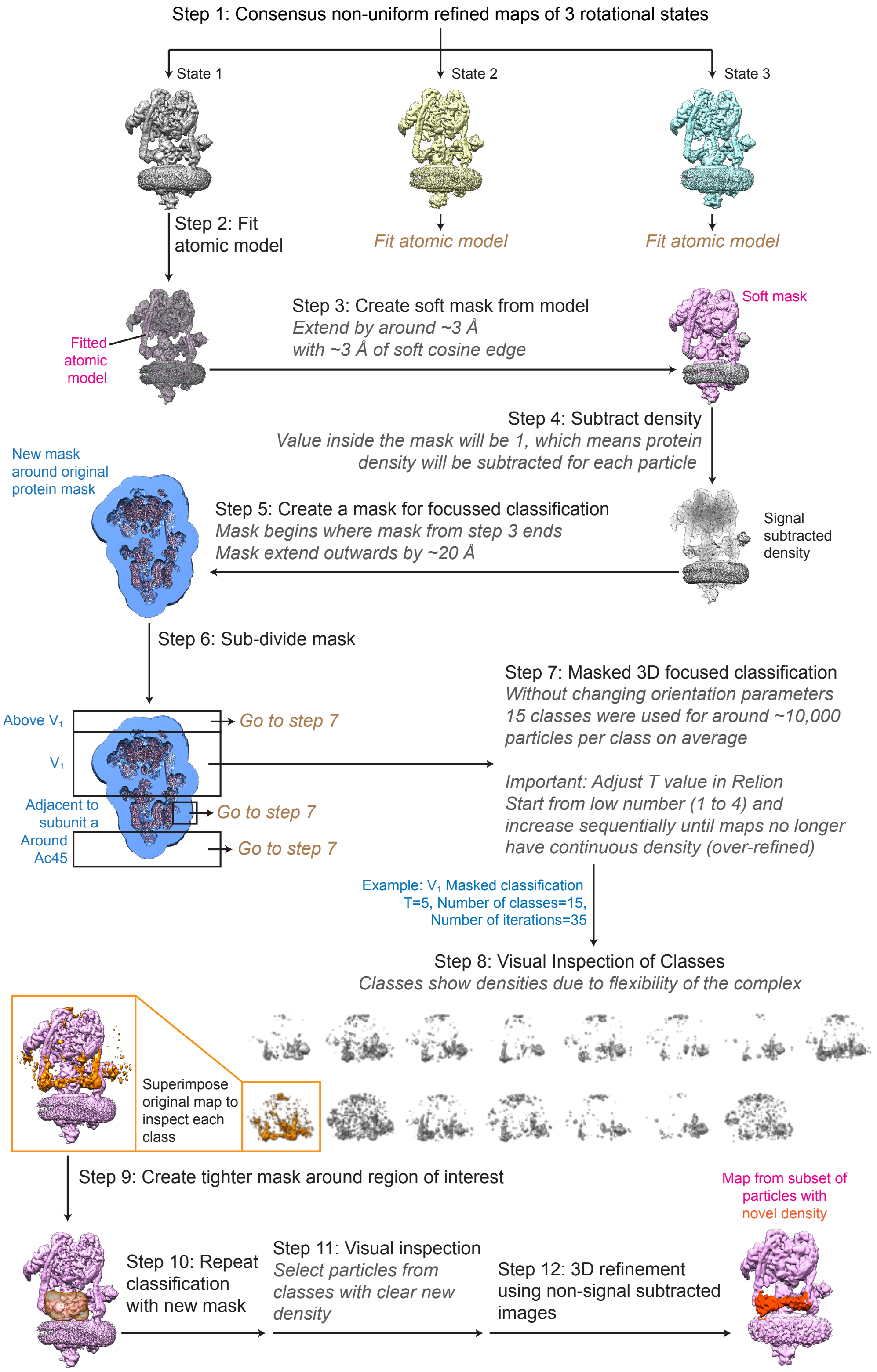


**Supplementary Figure 5. Exhaustive focused classification strategy for finding sub-stoichiometric proteins bound to V-ATPase.** The process is show for rotational State 1 as an example,but was applied to all rotational states.

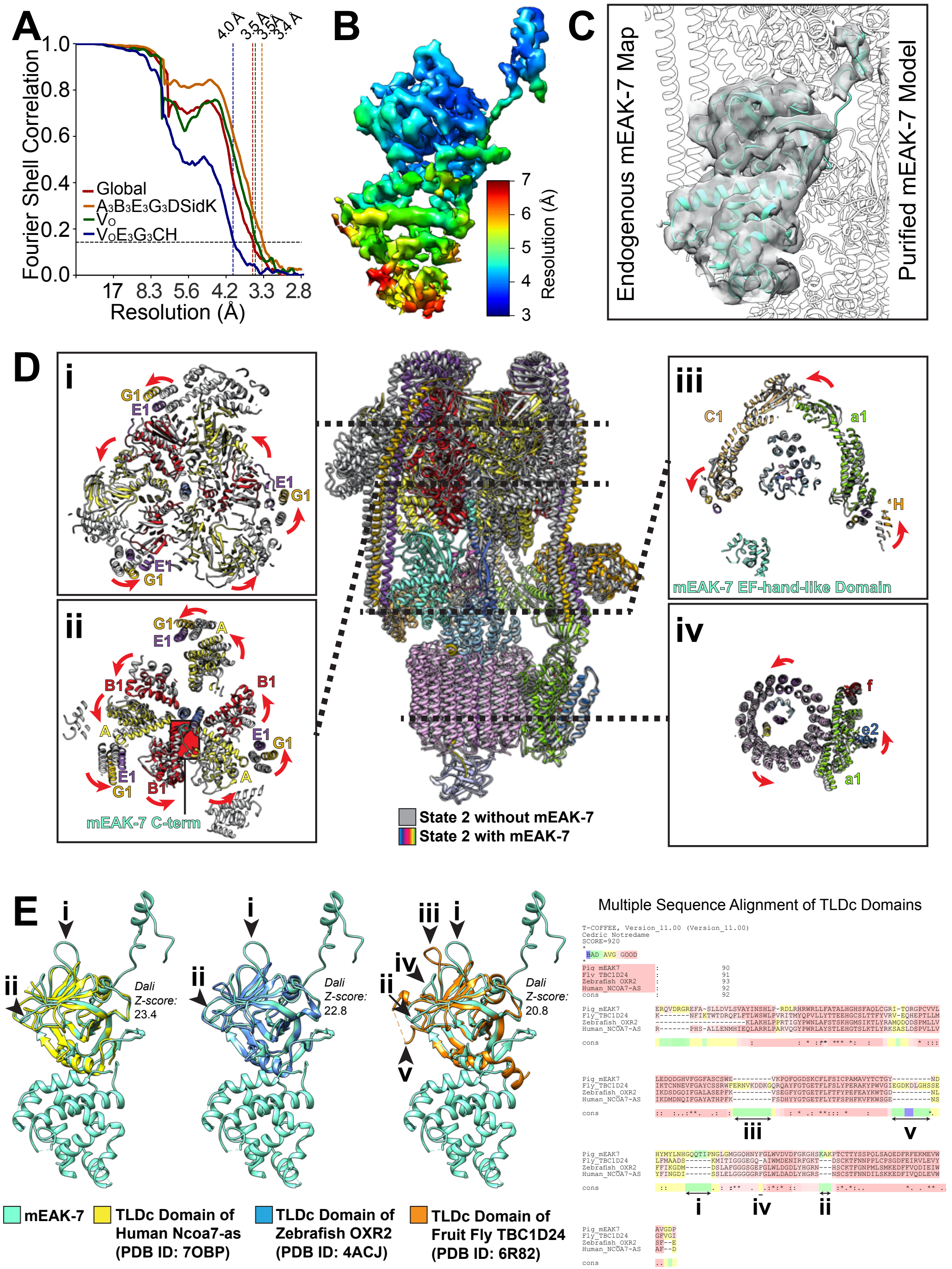


**Supplementary Figure 6. mEAK-7 binding to V-ATPase. A,** Fourier shell correlation curve following a gold-standard refinement and correction for masking for the map of V-ATPase bound to mEAK-7. **B,** Local resolution for mEAK-7 in the cryoEM map of mEAK-7 bound to V-ATPase. **C,** Comparison of the map for endogenous mEAK-7 (*grey surface*) and atomic model for recombinant mEAK-7 (*cyan ribbons*) bound to V-ATPase shows that the two structures are equivalent. **D,** Conformational changes to V-ATPase upon mEAK-7 binding. **E,**

Comparison of known structures of TLDc domains (left) and multiple sequence alignment (*right*) shows numerous insertions (numbered with Roman numerals for each structure).

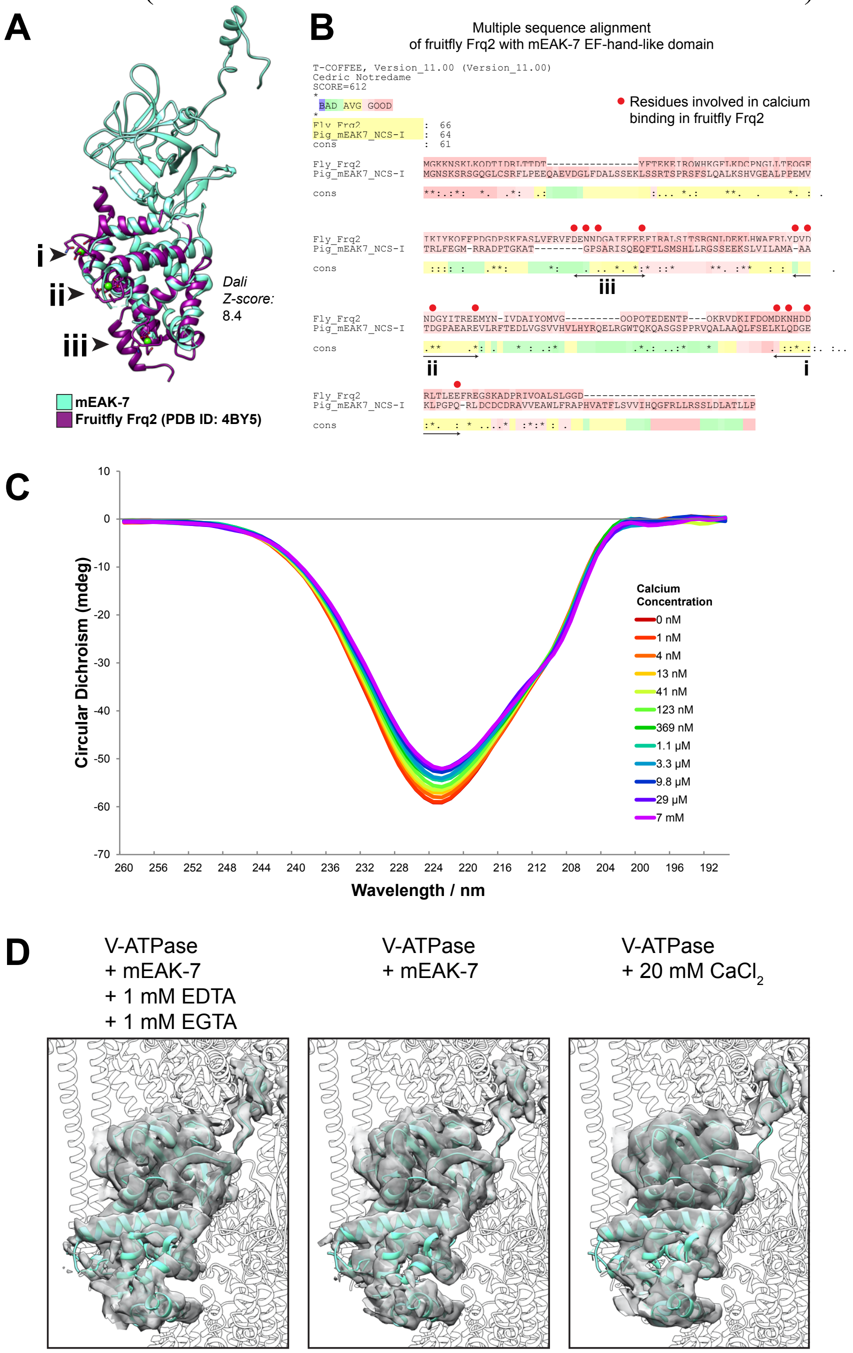


**Supplementary Figure 7. The EF-hand-like domain in mEAK-7 does not bind calcium. A,** Structural comparison of mEAK-7 and the EF-hand-like domain in fruit fly Frq2. The three calcium binding sites are numbered with roman numerals. **B,** Sequence comparison of mEAK-7 and the NCS-I domain in fruit fly Frq2, with red dots showing the residues in Frq2 that are involved in calcium binding and Roman numerals corresponding to the three calcium binding sites indicated in (A). Note that these residues are not well conserved in mEAK-7. **C,** Circular dichroism spectroscopy of mEAK-7 with varying concentrations of calcium. **D,** Experimental CryoEM maps (*grey surfaces*) for mEAK-7 bound to V-ATPase with the chelators EGTA and

EDTA (*left*), no chelator and no calcium (*middle*), and 20 mM calcium (*right*). The atomic model for mEAK-7 (*cyan ribbons*) in the absence of chelator or calcium is shown fitted into each map.

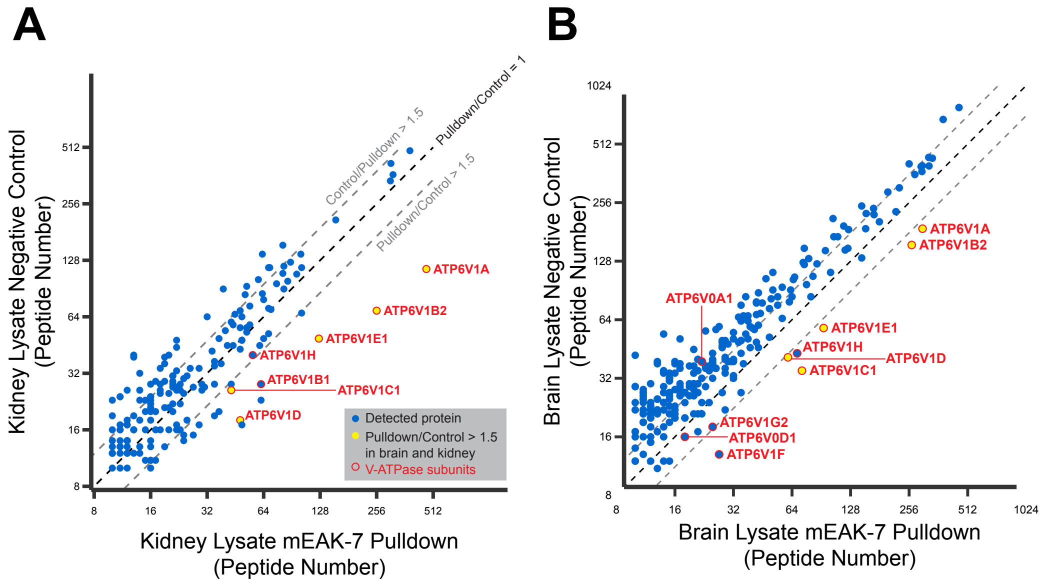


**Supplementary Figure 8. Mass spectrometry identification of proteins that bind mEAK-7.** Proteins that bind recombinant 3×FLAG-mEAK-7 upon incubation with kidney (**A**) and brain (**B**) lysate were identified and quantified approximately by the number of peptides found for each protein (*x-axis in each plot*). In a control experiment, proteins were quantified with an equivalent experiment that excluded 3×FLAG-mEAK-7 (*y-axis in each plot*).
